# Binding of CEP152 to PLK4 stimulates kinase activity to promote centriole assembly

**DOI:** 10.1101/2024.11.12.623205

**Authors:** Hazal Kübra Gürkaşlar, Ingrid Hoffmann

**Affiliations:** Cell Cycle Control and Carcinogenesis, D345, German Cancer Research Center, DKFZ, Im Neuenheimer Feld 242, 69120 Heidelberg, Germany; Faculty of Biosciences, Heidelberg University, 69120 Heidelberg, Germany

## Abstract

Centriole duplication is regulated by polo-like kinase 4 (PLK4) and several conserved initiator proteins. The precise timing and regulation of PLK4 activation are critical for ensuring that centriole duplication occurs only once per cell cycle. While significant progress has been made in understanding how PLK4 is activated, many aspects remain unclear. Here, we show how CEP152 contributes to the activation of PLK4. We utilize human cell lines that have been genetically engineered to rapidly degrade CEP152. Upon degradation of CEP152, localization of PLK4 at the proximal end of the centriole is disrupted. We show that binding of CEP152 N-terminal part to PLK4 increases phosphorylation and kinase activation. CEP152 controls the localization and levels of phosphorylated PLK4 at the proximal end of the centriole. CEP152 can phosphorylate and activate PLK4 both *in vitro* and *in vivo* which might stabilize PLK4 dimer formation, thus allowing autophosphorylation. We propose that CEP152 activates PLK4 to ensure proper centriole duplication at the onset of S-phase.

## INTRODUCTION

Centrioles are microtubule-based structures that are used to assemble both centrosomes and cilia/flagella. They recruit a surrounding proteinaceous pericentriolar material, PCM, to build the centrosome that forms the poles of the mitotic spindle during cell division (Nigg and Holland, 2018). Centrioles duplicate once during the cell cycle, specifically at the G1/S phase transition in that a new daughter centriole forms at the proximal end of the maternal centriole (Nigg and Holland, 2018). Centriole biogenesis must be tightly regulated to maintain proper centrosome number, as abnormalities can lead to chromosome missegregation and aneuploidy and have been linked to cancer (Nigg and Holland, 2018) (Banterle and Gonczy, 2017). Procentriole formation starts with the assembly of a cartwheel structure that is catalyzed by the conserved polo-like kinase 4, PLK4 (Nigg and Holland, 2018). Several scaffolding components CEP192/CEP152/CEP63/CEP57 ensure the presence of PLK4 at the proximal end of the maternal centriole (Cizmecioglu et al., 2010) (Hatch et al., 2010) (Sonnen et al., 2013) (Kim et al., 2013). CEP192 and CEP152 form distinct complexes with PLK4 by binding to the conserved polo boxes (PB) 1 and 2 located at its C-terminus. Double depletion of CEP152 and CEP192 causes a lack of centriolar PLK4 and prevents centriole assembly (Sonnen et al., 2013) (Kim et al., 2013) (Scott et al., 2023). CEP152 associates with nascent centrioles following disengagement during the first G1 phase (Kim et al., 2013) (Park et al., 2014). To initiate centriole duplication, PLK4 phosphorylates STIL at the conserved STAN motif, promoting the recruitment of SAS-6 (Dzhindzhev et al., 2014) (Ohta et al., 2014) (Kratz et al., 2015) (Moyer et al., 2015) a critical factor for cartwheel assembly.

The regulation of PLK4 protein levels is achieved through ubiquitylation and subsequent degradation via the proteasome. Previous studies have shown that this process is partially controlled by the SCF β-TrCP/SLIMB E3 ubiquitin ligase complex (Cunha-Ferreira et al., 2009) (Rogers et al., 2009) that recognizes a conserved phospho-degron generated through homodimer-dependent trans-autophosphorylation of PLK4 (Guderian et al., 2010) (Holland et al., 2010) (Cunha-Ferreira et al., 2013) (Klebba et al., 2013). CRL4^DCAF1^, which ubiquitylates PLK4 by binding to the conserved PB1-PB2 domain, is a complementary way to control PLK4 abundance to prevent centriole overduplication (Grossmann et al., 2024).

PLK4 phosphorylation is independent on the presence of STIL/SAS-6 and is already detectable before STIL/SAS-6 loading into a single site around the mother centriole (Yamamoto and Kitagawa, 2019). It will be important to assess how its activity and phosphorylation state are regulated during the cell cycle. PLK4 uniquely forms homodimers through PB1 and PB2 domains (Slevin et al., 2012) that allow trans-autophosphorylation (Guderian et al., 2010) (Holland et al., 2010) (Cunha-Ferreira et al., 2013) (Klebba et al., 2013). CEP152 is a strong binding partner of the PB1-PB2 of PLK4 and located at the centrosome prior to PLK4 loading. Thus, we analyzed whether PLK4 could be directly activated by binding to CEP152.

In this study, we genetically engineered human cells for fast and controllable degradation of CEP152 to better understand its dynamics and the consequences of its insufficiency on PLK4 localization and function. We find that CEP152 degradation does not affect total PLK4 levels at the centrosome but perturbs PLK4 localization at the proximal end of the mother centriole, resulting in a scattered localization of PLK4 at the centriole. We show that CEP152 co-localizes with active PLK4 at the proximal end of the mother centriole. Furthermore, we unravel the requirement of CEP152 in phosphorylation and activation of PLK4.

## RESULTS AND DISCUSSION

### Removal of CEP152 by auxin-dependent fast degradation

To gain more insight into the function of CEP152 in the localization and activation of PLK4 we removed CEP152 from cells. Depletion using shRNA or siRNA is not well controllable and might affect centriolar structure which is the case when targeting proteins involved in centriole duplication. In addition, rapid siRNA depletion during specific cell cycle phases is difficult. To circumvent this problem, we generated two cell lines, HeLa^CEP152-AID^ (Fig. 1) and U2OS^CEP152-AID^ (Fig. S1), in which CEP152 is rapidly degraded by an auxin-inducible degradation system (AID). This method can conditionally induce the degradation of CEP152 by the proteasome (Fig. 1A) simply by adding indole-3-acetic acid (IAA) to the cell culture medium (Nishimura and Fukagawa, 2017; Nishimura et al., 2009). All endogenous CEP152 alleles were inactivated using CRISPR/Cas9 and CEP152-AID was randomly integrated into the genome (Fig. 1B). Blasticidin-resistant clones used in this study were sequenced (Fig. 1B and S1A) to ensure the absence of endogenous CEP152 (Fig. 1C). CEP152-AID was degraded within 30 min after IAA addition (Fig. 1D) and restored within 2 h of IAA washout (Fig. 1E). Similarly, immunofluorescence signal showed a reduction of the CEP152 signal at the centrosome in response to IAA-treatment (Fig. 1F, S1E). A similar effect of CEP152-AID degradation was observed in U2OS cells (Fig. S1B-D). After 2 h of IAA depletion, we used ultrastructure expansion microscopy, U-ExM (Gambarotto et al., 2019) (Gambarotto et al., 2021) and stained centrioles with acetylated tubulin. We find that centriole length and diameter were not altered suggesting that its function was not impaired (Fig. 1G, S1F). On the contrary prolonged degradation of CEP152 for 24 to 72 h leads to a time-dependent impairment of centriole structure integrity resulting in short, narrow and broken centrioles (Fig. 1H). Short-term degradation of CEP152 (2 h) did not impair centriole number (Fig. 1I) compared to long-term treatment (72 h) implying that no functional defects accumulated. Together, we find that HeLa^CEP152-AID^ and U2OS^CEP152-AID^ are an ideal model to deplete CEP152 rapidly from centrosomal populations and study the effects of acute CEP152 deficiency on centriole assembly and function in cell lines.

**Figure 1:**
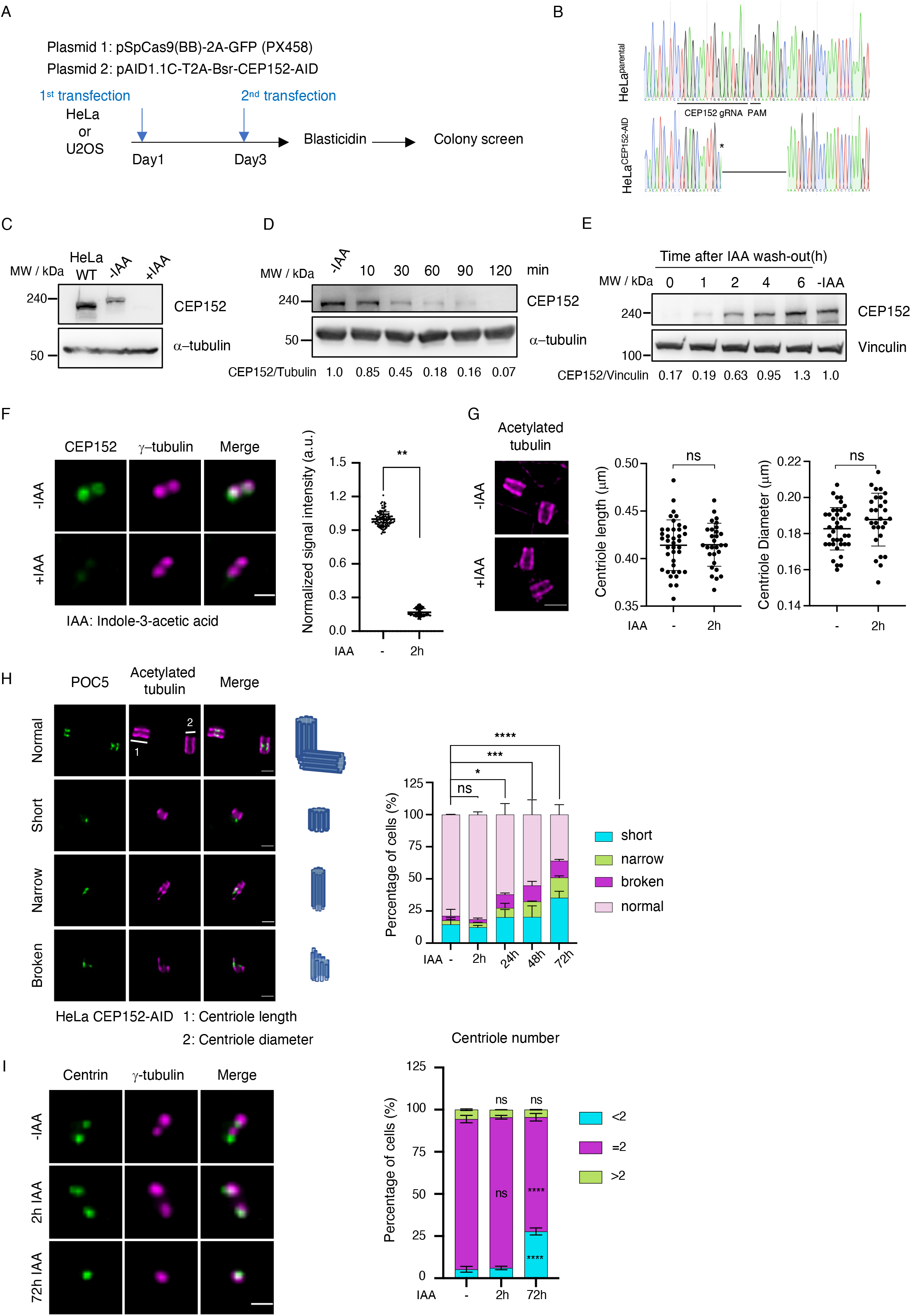
Fast degradation of CEP152 fused to AID. **(A)** Scheme of transfections and selection strategy to generate HeLa and U2OS cell lines expressing CEP152 fused to an AID. Sequencing result confirming the inactivation of CEP152 alleles in HeLa CEP152-AID cells by a deletion mutation. * point mutation. **(C)** HeLa CEP152-AID cells were treated with or without IAA for 2 h. Cells were lysed and analyzed by Western blotting. **(D)** HeLa CEP152-AID cells treated with or without IAA for indicated time points. Cells were lysed and analyzed by Western blotting. **(E)** HeLa CEP152-AID cells were treated with or without IAA for 2 h. IAA was washed-out for indicated time points and cells were lysed for Western blot analysis. The ratio of CEP152 / a-tubulin is indicated below the blot **(F)** Immunofluorescence staining of HeLa CEP152-AID cells after fixation depicts fast degradation of CEP152 upon 2 h IAA treatment. Scale bar: 1 μm. Values were normalized to the non-treated control and individual values were presented in the graph with ± SD. n = 2, ** < 0.01. **(G)** HeLa CEP152-AID cells were treated with IAA for 2 h and processed for U-ExM. Gels were stained with acetylated tubulin and the length and the diameter of centrioles were measured. The measured numbers were divided by expansion factor of the gels to obtain biological values. Scale bar: 1 μm, biological scale: 0.6 μm. Graphs represent the quantification of centriole length and diameter, n = 2, ns > 0.05. **(H)** Representative U-ExM images of HeLa CEP152-AID cells treated with IAA for the indicated times. Centriole schemes depicting approximate centriole orientation are included (left). Quantification graph shows the accumulation of defects on centriole structure in HeLa CEP152-AID cells treated with IAA for indicated time points with +SD. n = 2. * = 0.05, *** < 0.001, **** < 0.0001, scale bar: 1 μm, biological scale: 0.25 μm. (right). **(I)** HeLa CEP152-AID cells treated with IAA for 2 h and 72 h, fixed and stained with the indicated antibodies. Centriole number is counted by using centrin signal and results are shown in the graph with ± SD. ns > 0.05, ** < 0.01, **** < 0.0001. Scale bar: 1 μm.

### Acute removal of CEP152 disrupts the localization of PLK4 at the centrosome

To understand how the loss of CEP152 would affect PLK4 protein levels at the centrosome, we treated HeLa^CEP152-AID^ cells with IAA for 2 h to induce rapid degradation of CEP152. Acute removal of CEP152 did not alter PLK4 protein levels at the centrosome (Fig. 2A). Interestingly, we observed that upon fast degradation of CEP152, PLK4 lost its proximal end localization where the new centriole is built leading to a scattered pattern of PLK4 signal around the centriole (Fig. 2B). This suggests that the presence of CEP152 ensures proper localization of PLK4 at the proximal end of the procentriole. CEP63 and CEP152 form a hetero-tetrameric a-helical bundle, which is essential for their cooperative function in centrosomes (Kim et al., 2019). Binding of CEP63 to CEP152 forms a cylindrical platform that influences the level of PLK4 recruitment to the centrosome (Wei et al., 2020). We studied the effect on PLK4 loading in the absence of both CEP152 and CEP63 in cells where CEP63 was depleted in HeLa^CEP152-AID^ cells that were additionally treated with IAA for 2 h to degrade CEP152. Centriolar structure was not impaired upon short term degradation of CEP152 and siRNA depletion of CEP63 (Fig. S2A). As shown in Fig. 2C, degradation of CEP152 did not alter PLK4 levels at the centrosome while depletion of CEP63 slightly reduced the signal of PLK4 at the centrosome. Simultaneous depletion of CEP63 and CEP152 degradation in HeLa^CEP152-AID^ cells, however, lead to a three-fold reduction of the centrosomal PLK4 signal. Similar results were obtained upon depletion of CEP63 in U2OS^CEP152-AID^ cells treated with IAA for rapid CEP152 degradation (Fig. S2B). We find that PLK4 exists in a complex with CEP63 after co-immunoprecipitation. Although binding between PLK4 and CEP63 was reduced in the absence of CEP152, we anticipate that the interaction between PLK4 and CEP63 is either not direct or that CEP152 and CEP63 collaborate to regulate PLK4 levels at the centrosome (Fig. 2D). Together, our data show that CEP63 alone might not be able to recruit PLK4 to the centrosome but in cooperation with CEP152 which directly binds PB1 and PB2 of PLK4 (Kim et al., 2019).

**Figure 2:**
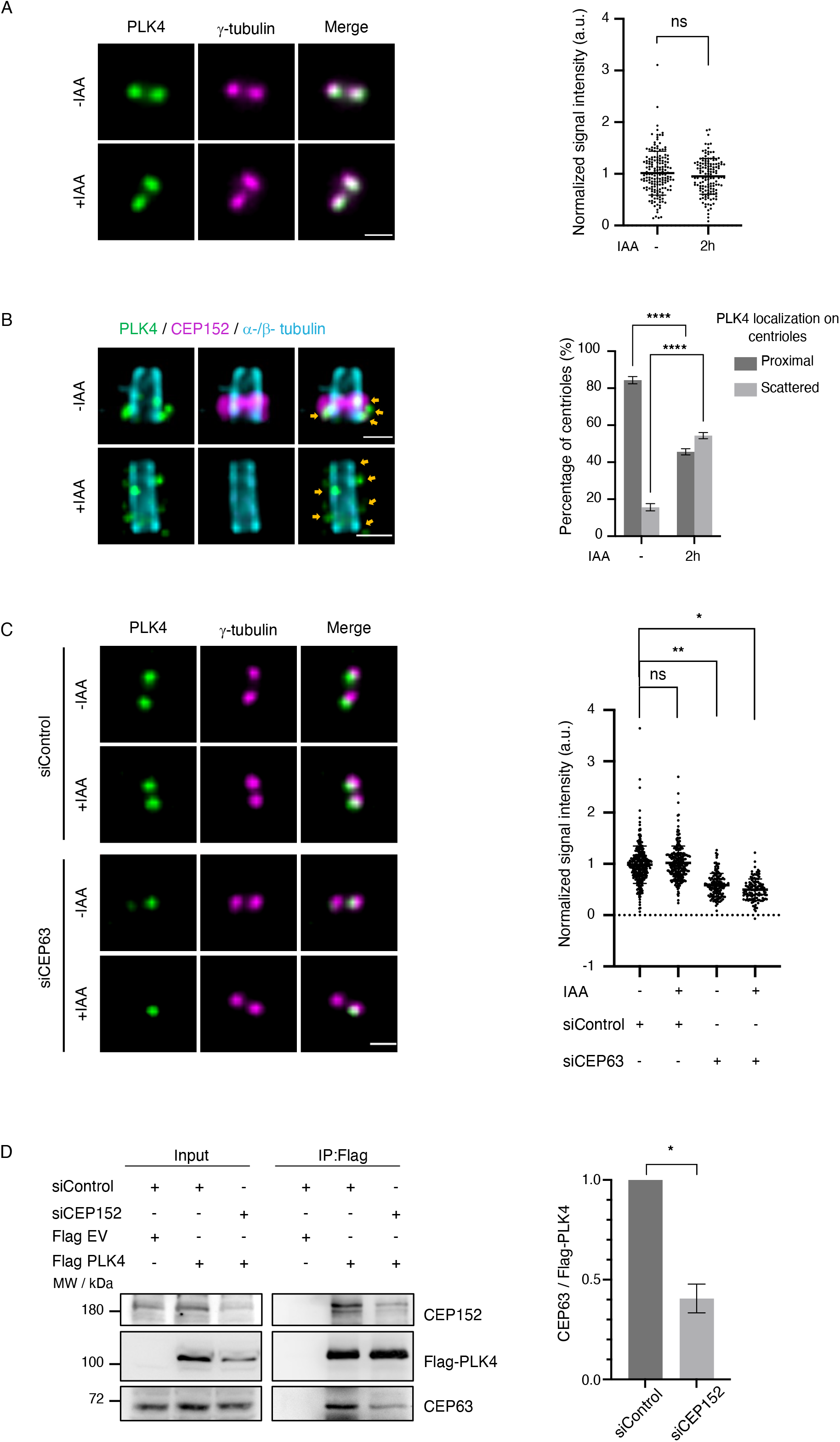
Fast degradation of CEP152 disrupts PLK4 localization at the centrosome. **(A)** HeLa CEP152-AID cells were treated with or without IAA for 2 h and cells were then fixed for immunofluorescence analysis. Scale bar: 1 μm. PLK4 signal was normalized to the non-treated control and values were shown in the graph with ± SD. ns > 0.05. **(B)** HeLa CEP152-AID cells were treated with IAA for 0 h or 2 h, fixed and stained with the indicated antibodies for U-ExM analysis. Arrows indicate PLK4 foci. Quantification graph of U-ExM images shows the percentage of the cells with either proximal or scattered PLK4 distribution at the centrosomes after rapid degradation of CEP152. n = 2, Scale bar: 1 μm, biological scale: 0.2 μm. **(C)** HeLa CEP152-AID cells transfected with control (siGL2) and CEP63 siRNAs for 48 h were treated or not with IAA for 2 h. Samples were stained with indicated antibodies. Images show the change in PLK4 levels at the centrosome. Scale bar: 1 μm. PLK4 signal was normalized to the non-treated control and values were shown in the graph with ± SD n = 3. * = 0.05, ** < 0.01. **(D)** HEK293T cells were transfected with siRNAs against GL2 (control) and CEP152. Cells were then transfected with empty Flag vector or Flag PLK4 for 48h. Cells were harvested 72 h after the first siRNA transfections for immunoprecipitation and Western Blot analysis using the indicated antibodies. n = 3, * = 0.05.

### CEP152 promotes PLK4 autophosphorylation and activation

Our data suggests that CEP152 is required for proximal end localization of PLK4, where the new centriole is built (Fig. 2B). We therefore set out to investigate what function CEP152 might play in the regulation of the activity and phosphorylation state of PLK4. PLK4 negatively regulates its own stability through multiple trans-autophosphorylation at a degron motif in the L1 region, an unstructured linker following the PLK4 kinase domain (Holland et al., 2010) (Scott et al., 2023) (Lopes et al., 2015). We realized, however, that a full depletion of CEP152 is difficult to achieve upon degradation of CEP152 in HeLa^CEP152-AID^ cells as IAA-treatment did not lead to a complete removal of CEP152. To study a possible role of CEP152 in the activation of PLK4, we need to assure that CEP152 is not detectable on centrosomes in HeLa cells. To achieve this, we used HeLa^CEP152-AID^ cells to degrade CEP152 and treated them additionally with CEP152 siRNA for 48 h to further reduce CEP152 levels (Fig. 3A). We only analyzed those cells where the CEP152 signal was not present at centrosomes. The structure of the centrioles was not impaired following the treatment (Fig. S3A). To monitor PLK4 autophosphorylation, we used an anti-PLK4 antibody raised against phosphorylated serine 305, p-S305 PLK4 (Park et al., 2014). S305 is located within the multiphosphodegron of the PLK4 L1 domain (Sillibourne et al., 2010) (Park et al., 2014). This antibody monitors autophosphorylation of PLK4 which is a consequence of kinase activation (Sillibourne et al., 2010). p-S305 PLK4 signals start to be detected and simultaneously increase before the loading of STIL/SAS-6 (Yamamoto and Kitagawa, 2019). While total PLK4 levels at the centrosome were not impaired in the absence of CEP152 in HeLa^CEP152-AID^ cells, the p-S305 signal was markedly decreased suggesting that CEP152 might regulate PLK4 phosphorylation and activity (Fig. 3B). Similar results were also obtained in U2OS^CEP152-AID^ cells (Fig. S3B). The p-S305 PLK4 signal clearly decreased at the daughter centriole in the absence of CEP152, suggesting that p-S305 PLK4 is removed first from the daughter centriole upon degradation of CEP152 (Fig. S3C).

**Figure 3:**
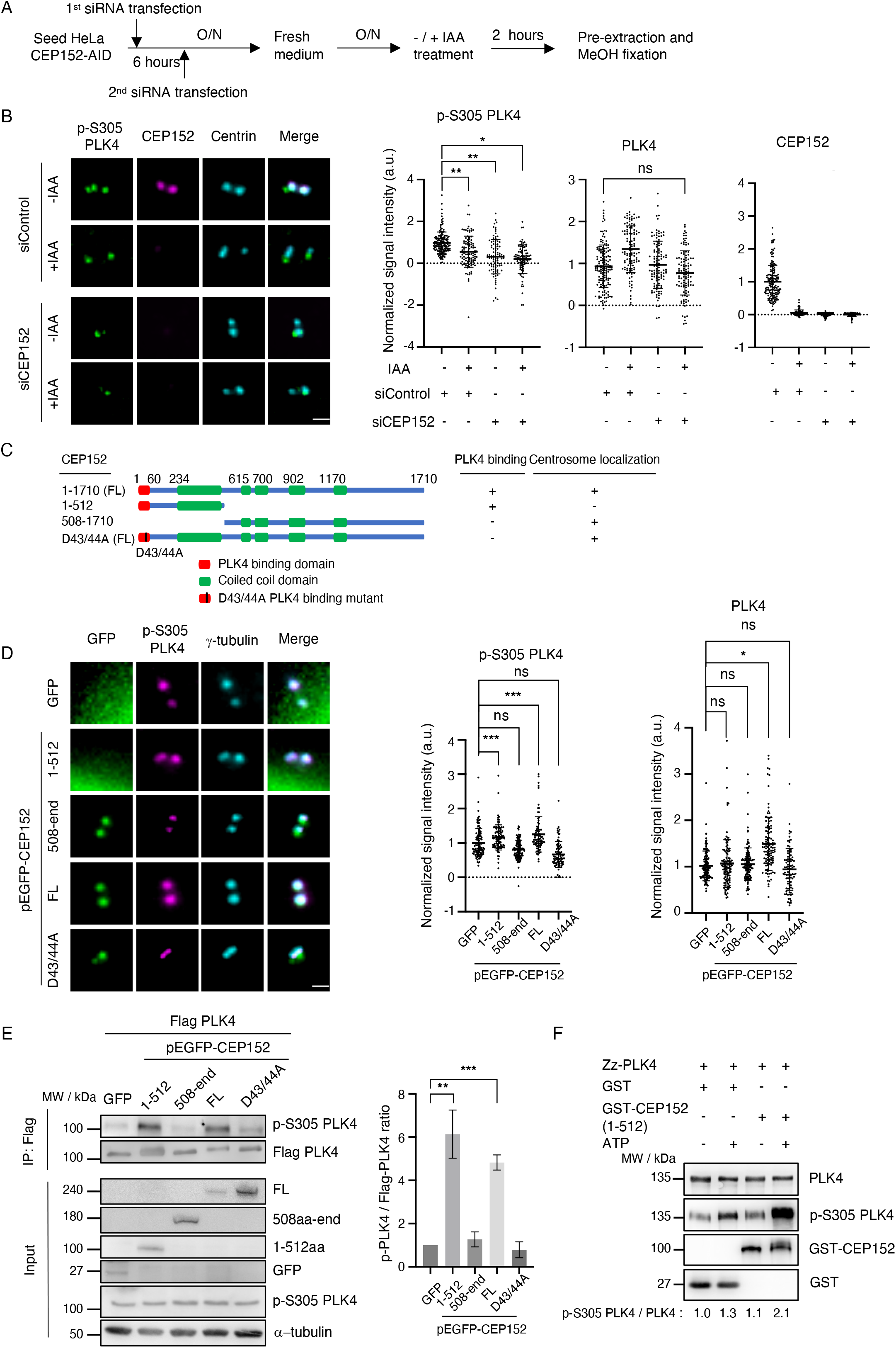
PLK4 phosphorylation at S305 is dependent on CEP152 binding. **(A)** Experimental work-flow to yield non-detectable CEP152 signals at centrosomes in HeLa CEP152-AID cells. **(B)** HeLa CEP152-AID cells were transfected with control or CEP152 siRNAs for 48 h and treated with - / + IAA for 2 h. Samples were fixed as in **A**. and stained. Scale bar: 1 μm. p-S305 PLK4 and PLK4 signals were quantified in cells with no detectable CEP152 and normalized to the control after background signal subtraction. Graphs show individual values n = 2, ns > 0.05, * = 0.05, ** < 0.01. **(C)** Schematic representation of GFP tagged full length (FL) CEP152, CEP152 1-512aa (N-term), 508-1710aa (C-term) and CEP152^D43/44A^ constructs and their ability to bind PLK4 and to localize at the centrosome. **(D)** HeLa WT cells were transfected with pEGFP-CEP152 constructs or pEGFP for 36 h. Cells were fixed after pre-extraction for immunofluorescence analysis. Scale bar: 1 μm. n = 3, ns > 0.05, * = 0.05, *** < 0.001. **(E)** HEK293T cells were co-transfected with Flag-PLK4 and different pEGFP-CEP152 constructs shown in **C** for 36 h and then harvested for co-immunoprecipitation and Western Blot analysis. Graph shows the ratio of p-S305 PLK4 / Flag-PLK4 with ± SD. n = 3, * = 0.05, ** < 0.01. **(F)** GST-CEP152 (1-512aa) or GST alone was incubated with Zz-PLK4-His in the presence and the absence of 0.5 mM ATP for *in vitro* kinase assay and then analyzed by Western blotting.

PLK4 and CEP152 interact both *in vitro* and *in vivo* in cells (Cizmecioglu et al., 2010) (Hatch et al., 2010). To examine whether CEP152 interaction with PLK4 would increase PLK4 autophosphorylation at S305, we used a binding mutant or truncated versions of CEP152 that either interact with PLK4 without localizing to the centrosome or vice versa localized at the centrosome without interacting with PLK4 (Cizmecioglu et al., 2010) (Fig. 3C). We found that expression of full-length CEP152 significantly increased p-S305 PLK4 levels at the centrosome (Fig. 3D). On the contrary, a CEP152 (D43A, D44A) double mutant that exhibited a significantly compromised ability to bind to the full-length PLK4 (Kim et al., 2013) did not cause activation of PLK4. Binding of an N-terminal CEP152 fragment was sufficient to induce phosphorylation of PLK4. Activation of PLK4 by CEP152 was not dependent on CEP152 localization to the centrosome as a C-terminal CEP152 fragment that localizes to the centrosome did not lead to phosphorylation of PLK4 at S305 (Fig. 3D). Phosphorylation of PLK4 on S305 by CEP152 could also be confirmed biochemically by performing co-immunoprecipitation experiments of PLK4 and CEP152 (Fig. 3E). To show that CEP152 promotes PLK4 autophosphorylation directly *in vitro*, we performed *in vitro* kinase assays with recombinant Zz-PLK4 and the recombinant N-terminal fragment of CEP152, CEP152 1-512, that contains the PLK4 binding site (Cizmecioglu et al., 2010). We observed a low activity of recombinant PLK4 that is most likely caused by dimerization and trans-autophosphorylation (Cunha-Ferreira et al., 2013). Interestingly, PLK4 kinase activity markedly increased upon binding of PLK4 to the N-terminal CEP152 fragment in the presence of ATP (Fig. 3F) as monitored by a clear increase in the p-S305 PLK4 signal. It is conceivable, that the low activity after PLK4 dimerization is not sufficient to initiate centriole duplication and that further binding of the PLK4 dimer to CEP152 is required to reach a threshold activity of PLK4 in order to start this process.

### CEP152 binding to PLK4 allows PLK4 dimerization and activation

Monomeric PLK4 is autoinhibited. Once PLK4 dimerizes, this autoinhibition is relieved, allowing for increased catalytic activity thus facilitating further autophosphorylation. PB1 and PB2 domains provide an intramolecular homodimerization platform to regulate PLK4 trans-autophosphorylation (Klebba et al., 2015) and also interact directly with CEP152 (Cizmecioglu et al., 2010) (Hatch et al., 2010) and CEP192 (Kim et al., 2013). As both dimerization of PLK4 and CEP152 binding involve PB1 and PB2, we checked whether CEP152 might affect PLK4 dimerization in human cells. If this is correct, dimerization of PLK4 should be reduced in the absence of CEP152. We find that upon depletion of CEP152, homodimerization of PLK4 was reduced which might imply that binding of CEP152 to PLK4 stabilizes dimerization and leads to activation of PLK4 but does not affect PLK4 protein levels at the centrosome (Fig. 2A). In fact, we find that depletion of CEP152 leads to a decrease in p-S305 PLK4 levels suggesting that dimerization is necessary for activation of PLK4 through phosphorylation of S305 (Fig. 4A). CEP152 competes with another scaffold protein, CEP192, for binding to PB1 and PB2 of PLK4 (Kim et al., 2013). We asked whether expression of CEP192 would affect PLK4 activation by competing with CEP152 for PLK4 binding thus impairing PLK4 activation. We therefore expressed increasing amounts of CEP192 and checked for changes in binding between CEP152 and PLK4. As expected, we find that increasing amounts of CEP192 lead to a stepwise decrease in CEP152 binding to PLK4. Moreover, this decrease in CEP152 binding led to a decrease in p-S305 PLK4 levels (Fig. 4B) as dimerization and activation of PLK4 is impaired. Together, these results suggest that binding of CEP152 to PLK4 stabilizes PLK4 dimerization leading to autophosphorylation. (Klebba et al., 2015) reported that Asterless, the ortholog of CEP152 in *D. melanogaster* has two PLK4 binding sites and that the N-terminal PLK4 binding site also affects PLK4 homodimerization. Our studies now confirm that binding of the N-terminal domain of CEP152 to PLK4 leads to enhanced PLK4 kinase activity in human cells.

**Figure 4:**
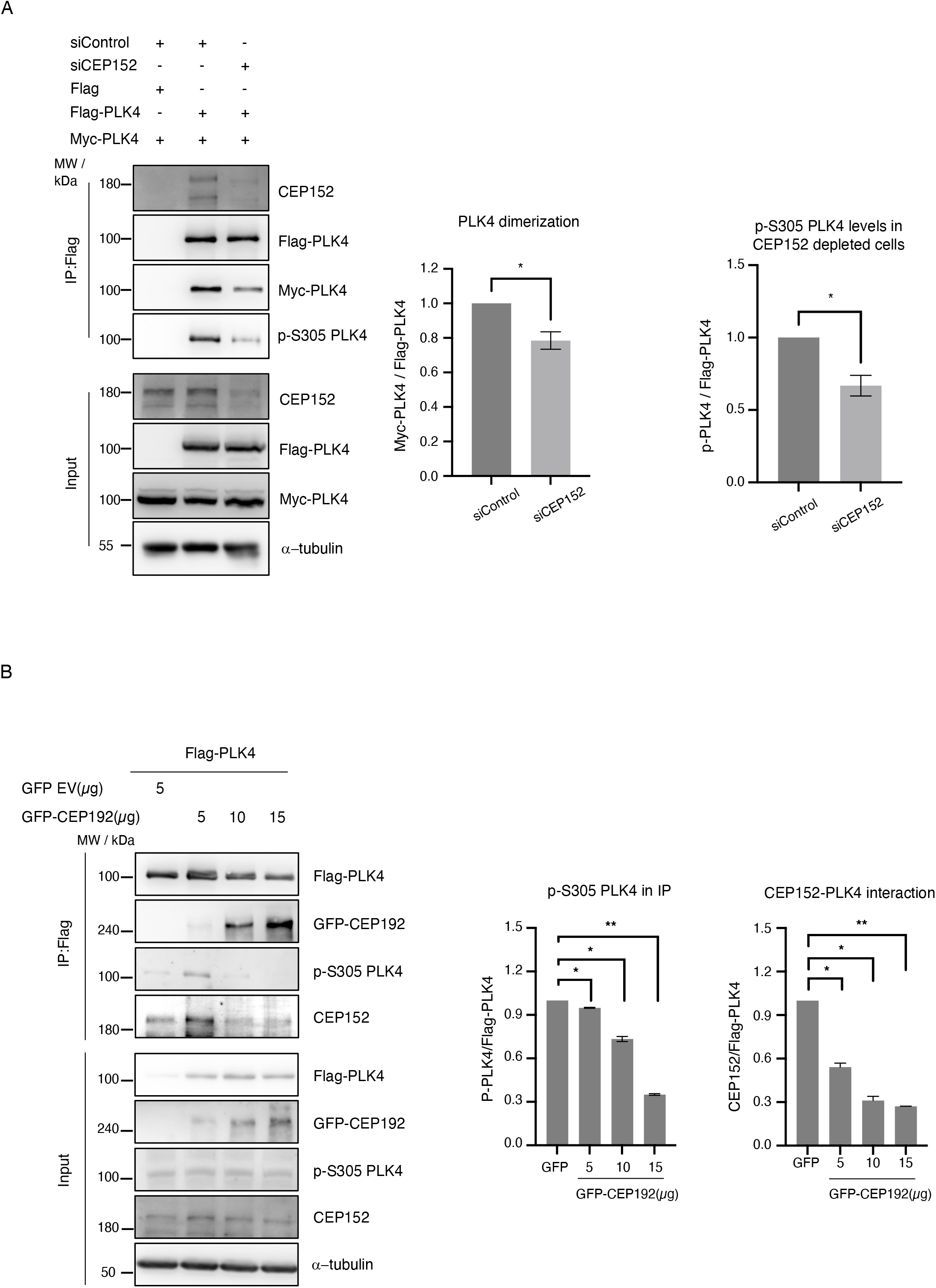
CEP152 regulates dimerization of PLK4. **(A)** HEK293T cells were transfected with siRNAs against GL2 (control) or CEP152. Cells were then co-transfected with empty Flag vector and Myc-PLK4 or Flag-PLK4 and Myc-PLK4 for 48h. Cells were harvested 72 h after the first siRNA transfections for immunoprecipitation (IP) analysis and immunostaining. Graph on the left shows the quantification. n = 3, * = 0.05. **(B)** HEK293T cells were co-transfected with Flag-PLK4 and pEGFP or increasing amounts of pEGFP-CEP192 for 36 h followed by Western blot detection. n = 3, * = 0.05, ** < 0.01.

### CEP152 promotes PLK4 activation at the proximal end of the procentriole

From our data it is conceivable that CEP152 regulates the activity and phosphorylation state of PLK4 at the site of procentriole formation. To analyze this in more detail, we first checked by U-ExM the co-localization of p-S305 PLK4 with CEP152 at the centriole in cells that were synchronized in S-phase. We used U2OS cells for this set of experiments, as the p-S305 PLK4 antibody did not work in U-ExM in HeLa cells presumably due to lower p-S305 PLK4 levels present in this cell line. CEP152 localizes as a continuous ring that encircles the base of the parental centriole but lacks a clear 9-fold organization (Sullenberger et al., 2023) (Scott et al., 2023). p-S305 PLK4 is primarily localized at the proximal end of the parental centriole where the new centriole is formed and where it co-localizes with CEP152 (Fig. 5A). Recently, it was shown that inactive PLK4 binds up to nine sites at the proximal end of parental centriole in HeLa cells (Scott et al., 2023). To examine whether CEP152 is involved in activation of PLK4 at the onset of S-phase, we used a protocol where U2OS^CEP152-AID^ cells were arrested in S-phase and then treated with centrinone to inhibit PLK4 kinase activity in the presence and absence of CEP152 (Fig. 5B). This would allow us to examine a change in the number of foci following kinase inhibition. Using this setup, we checked for the effect on PLK4 S305 phosphorylation upon fast degradation of CEP152 followed by co-staining for total PLK4 and p-S305 PLK4 to exactly monitor the localization of active PLK4. After centrinone-washout the p-S305 PLK4 signal was visible at 4 PLK4 sites. Upon rapid degradation of CEP152, the number of p-S305 PLK4 foci clearly decreased to 0-1 foci at parental centrioles, suggesting that PLK4 is inactive in the absence of CEP152 at the proximal end of the procentriole (Fig. 5C). Next, we compared the localization of p-S305 PLK4 in U2OS CEP152-AID cells with and without IAA treatment. Interestingly, the localization of p-S305 PLK4 was not restricted to the proximal end of centrioles. While 45% of centrioles exhibited only proximal-end localization of p-S305 PLK4, around 35% showed both proximal and distal-end localization. Thus, the majority of cells (∼85%) displayed proximal localization of p-S305 PLK4 (Fig. 5D). Strikingly, this number dropped to 30% in CEP152-degraded U2OS^CEP152-AID^ cells. As expected, nearly 50% of the cells showed no p-S305 PLK4 localization. However, the number of cells with distal-end-only localization of p-S305 PLK4 increased to 25% (Fig. 5D). This suggests that CEP152 is required to restrict p-S305 PLK4 to the proximal end, where centriole duplication is initiated. The distinct localization patterns of p-S305 PLK4 observed in this study raise important questions about the spatial regulation of PLK4 activity and its functional consequences. The increase in distal-only p-S305 PLK4 localization in the absence of CEP152 raises the possibility of deregulated centriole assembly, which could affect centriole integrity and function (Fig. 5D), suggesting that short-term degradation of CEP152 decreases the phosphorylation of PLK4 at S305. Finally, we analyzed SAS-6 loading, which is essential for cartwheel formation and proper centriole duplication, in response to degradation of CEP152 in S-phase arrested cells. We noticed that SAS-6 localization was clearly reduced in the absence of CEP152 (Fig. 5E). These results suggest that CEP152 is also crucial for the recruitment of SAS-6 to ensure that phosphorylation of STIL by PLK4 triggers the loading of SAS-6 onto both mother and daughter centrioles after they disengage at the end of mitosis.

**Figure 5:**
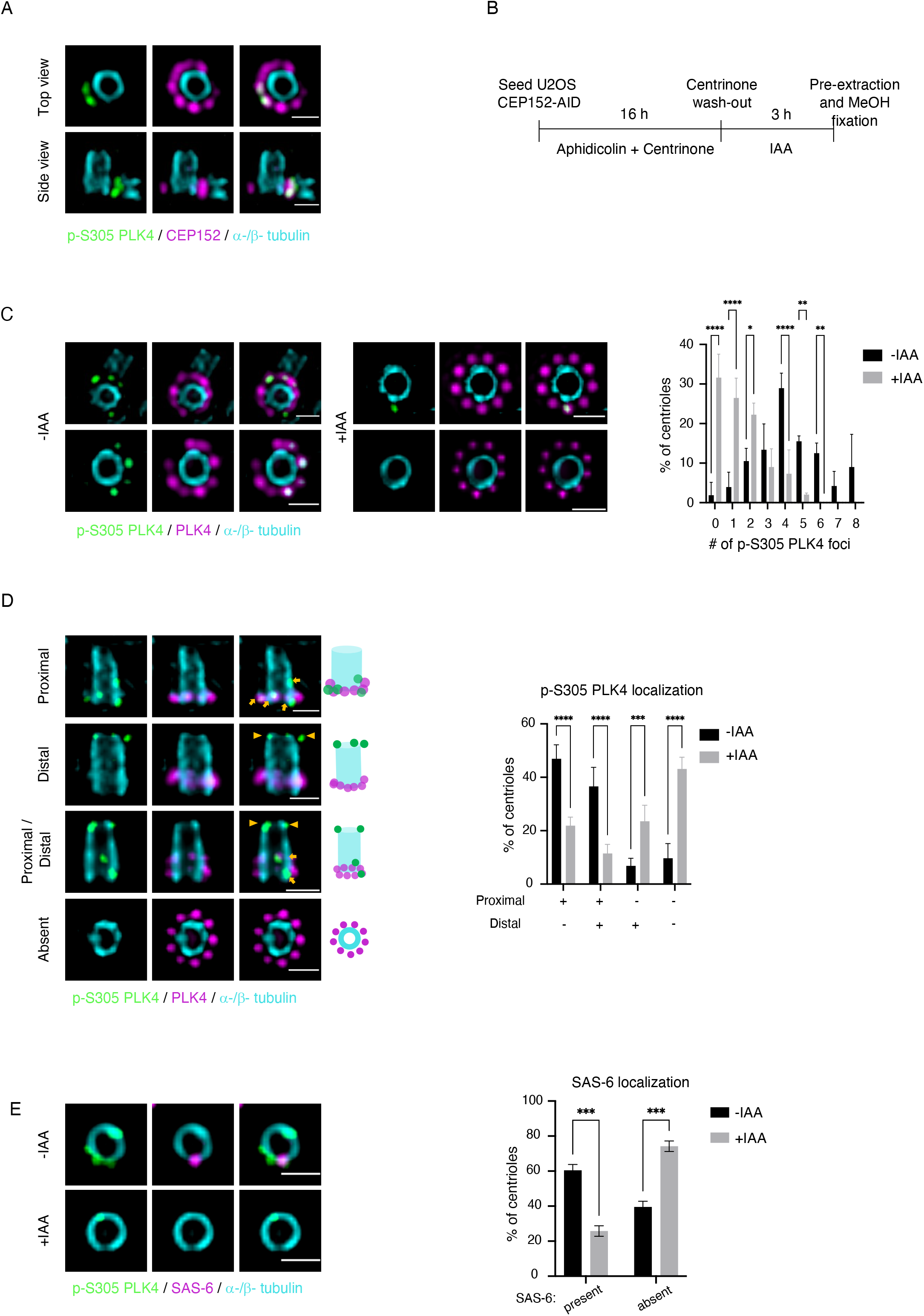
CEP152 regulates proximal end localization and levels of p-S305 PLK4 at the centriole. **(A)** U2OS WT cells were synchronized in early S-phase with 5 mg/ml Aphidicolin for 16 h and active PLK4 was enriched by treatment with 0.1 μM centrinone for 16 h and wash-out for 3 h. Fixed samples were used for U-ExM analysis. **(B)** Experimental workflow of S-phase synchronization, centrinone wash-out and IAA treatments of U2OS CEP152-AID cells. U2OS CEP152-AID cells were treated as indicated in **(A)** and fixed samples were used for U-ExM analysis. Gels were stained with the indicated antibodies in - / + IAA treated U2OS CEP152-AID cells. The number of p-S305 PLK4 foci was counted and the percentage of centrioles was shown in the graph. * shows p-S305 PLK4. **(D)** Representative U-ExM images of different p-S305 localizations in U2OS CEP152-AID cells after the protocol shown in (**B)**. Arrowheads and arrows mark distal and proximal localization of p-S305 PLK4, respectively. Graph shows the percentage of the centrioles with different p-S305 PLK4 localizations in - / + IAA treated U2OS CEP152-AID cells. **(E)** Representative U-ExM images of SAS-6 localization in - / + IAA treated U2OS CEP152-AID cells. Graph shows the percentage of centrioles with SAS-6 recruitment in - / + IAA treatments. Scale bars: 1 μm for 4-fold expansion images.

Based on our findings we propose that CEP152 is involved in the activation of PLK4 in S-phase by binding to the PB1-PB2 domains of PLK4, allowing PLK4 dimerization and activation through phosphorylation. A crucial unanswered question is how PLK4 kinase activity is temporally controlled to trigger centriole assembly. CEP152 is found at the centrosome throughout the cell cycle, with its levels fluctuating depending on the phase. In interphase, it is predominantly localized around centrioles, while during mitosis, its presence is reduced but still significant (Cizmecioglu et al., 2010) (Hatch et al., 2010). CEP152 associates with newly formed centrioles following their disengagement during the first G1 phase (Kim et al., 2013) (Park et al., 2014). In early G1, CEP152 localizes at the mother centriole and then repositions PLK4 to the outer edge of a newly forming CEP152 ring at the daughter centriole (Park et al., 2014). PLK4 is slightly activated through dimerization in the absence of CEP152 (Fig. 3F) but the level of kinase activity might not be enough to trigger centriole duplication in cells. Following repositioning, we propose that binding of CEP152 to PLK4 increases PLK4 kinase activity of the dimer at the proximal end of the centriole as this localization is lost in the absence of CEP152 (Fig. 5D). Apart from CEP152, STIL binding to PLK4 also stimulates its kinase activity by promoting self-phosphorylation at T170 located in the activation loop (T-loop) leading to increased activity. In cycling cells and in contrast to PLK4 phosphorylation by CEP152, STIL induced PLK4 phosphorylation leads to subsequent destruction of PLK4 (Moyer et al., 2015) while PLK4 protein levels remain unaltered in response to CEP152 binding. Our data provide new evidence on how PLK4 is regulated over time. We propose that CEP152 dependent activation of PLK4 occurs during early G1 to trigger STIL recruitment to centrioles by PLK4. Phosphorylation of STIL by active PLK4 then facilitates STIL/SAS-6 interaction and centriolar loading of SAS-6. The conserved TIM domain in STIL stimulates autophosphorylation and degradation of PLK4 (Ohta et al., 2018). It would be interesting to show how exactly CEP152 binding leads to activation of PLK4. One possibility is that CEP152 stabilizes dimerization of PLK4 by binding to PB1 and PB2. Alternatively, CEP152 might trigger a conformational change that allows more efficient kinase activation. Further investigation will shed light on the temporal activation of PLK4 to expand our understanding of centriole duplication.

## MATERIALS AND METHODS

### Cell culture, generation of stable cell line, transfection of plasmids and siRNAs

HeLa (ATCC CCL-2), U2OS (ATCC HTB-96), HEK293T (ACC 635; DSMZ, Braunschweig, Germany), HeLa CEP152-AID and U2OS CEP152-AID cells were grown in high-glucose DMEM (Life Technologies) supplemented with 10% fetal bovine serum (FBS) (Sigma-Aldrich) and 1 % penicillin-streptomycin (Sigma-Aldrich) at 37°C and 5% CO2.

HeLa CEP152-AID and U2OS CEP152-AID cells stably expressing AID-fused CEP152 were generated according to the protocol established by (Vasquez-Limeta et al., 2022). Briefly, wild-type HeLa and U2OS cells were co-transfected with pAID 1.1C-T2A-Bsr-containing gRNA resistant CEP152 to express AID-fused CEP152 and pX458 Cas9-containing CEP152 gRNA to knock-out endogenous CEP152, and they were selected with 2 and 5 μg/ml Blasticidin S HCl (Santa Cruz Biotechnology). Colonies were screened for CEP152 knock-out and CEP152-AID expression. The degradation of AID-fused CEP152 was induced by the addition of 500 μM Indole-3-acetic acid (IAA) for 2 h (Nishimura and Fukagawa, 2017; Nishimura et al., 2009). Transfections of HeLa and U2OS cells were performed with Lipofectamine 2000 and Lipofectamine RNAiMax (Life Technologies), using 2 μg of plasmid and 50 nM siRNA respectively, to transfect six-well plates according to the manufacturer’s recommendations. HEK293T cells were used to perform immunoprecipitation experiments and transfections were performed with Polyethylenimine (Polysciences) at a final concentration of 5 μg/ml using 30 μg plasmid DNA per 15 cm^2^ dish (2 × 10^7^ cells).

To analyze PLK4 phosphorylation, cells were synchronized at early S phase by using 5 μg/ml Aphidicolin (Biomol) and PLK4 was inhibited by adding 0.1 μM centrinone for 16 h. For the activation analysis of PLK4, synchronized cells were washed with phosphate buffered saline (PBS) 5 times and released into fresh -/+ IAA-containing medium for 3 h. For siRNA experiments, CEP63 and CEP152 siRNAs were transfected into U2OS CEP152-AID or HeLa CEP152-AID cells for 72 h and 48 h, respectively, and treated with -/+ IAA during the last 2-3 h. Cells were fixed and processed for immunofluorescence (IF) or expansion microscopy (U-ExM) analysis.

### Antibodies

Mouse anti-FLAG M2 (1:5000 in Western blotting, WB; F3165), mouse anti-a-tubulin (1:1000 WB; T5168), mouse anti-acetylated-tubulin (1:1000 U-ExM; 6-11B-1) and mouse anti-y-tubulin (1:2000 IF; GTU-88) were obtained from Sigma. Mouse anti-centrin-2 (1:500 IF; 20H5) from Merck, mouse anti-CEP152 (1:1000 WB and 1:2000 IF; P1285) from Invitrogen, mouse anti-CEP192 (1:500 WB; A302-324A), rabbit anti-STIL (1:1000 IF; A302-442A) from Bethyl, mouse anti-MYC (1:1000 WB; 9E10), rabbit anti-GST (1:1000 WB; Z-5) mouse anti-ninein (1:1000 IF; F5), mouse anti-SAS-6 (1:500 IF, 1:100 U-ExM) from Santa Cruz Biotechnology, and rabbit anti-pericentrin (1:2000 IF; ab4448) from Abcam were used in different techniques. Mouse anti-centrobin (1:10000 IF), rabbit anti-a-tubulin (1:500 U-ExM), rabbit anti-?-tubulin (1:500 U-ExM) were obtained from Proteintech and Guinea Pig anti-a-tubulin (1:1500 U-ExM) and Guinea Pig anti-?-tubulin (1:1500 U-ExM) from ABCD antibodies. Rabbit anti-CEP152 (1:5000), rabbit anti-PLK4 (1:1000 IF, 1:500 U-ExM) and mouse anti-PLK4 (1:500 WB, 1:500 IF and 1:200 U-ExM) were used as in (Cizmecioglu et al., 2010). Rabbit anti-p-S305 PLK4 (Park et al., 2014) was a gift from Kyung Lee, NIH, USA. Secondary antibodies for IF and U-ExM were goat anti-guinea pig coupled to Alexa Flour 594, goat anti-mouse IgG, IgG_1_, IgG_2a_ or IgG_2b_, and goat anti-rabbit IgG coupled to Alexa Fluor 488, Alexa Fluor 594 and Alexa Flour 647 (1:500–1:1000; Molecular Probes). Secondary antibodies for Western blotting were peroxidase-conjugated donkey anti-rabbit (1:2000; Jackson Laboratories) and goat anti-mouse (1:2000; Novus).

### siRNAs and gRNAs

siRNAs used in this study were directed against the following sequences: firefly luciferase (GL2, control), 5′-AACGUACGCGGAAUACUUCGA-3′; CEP152 Oligo, 5′-GGAUCCAACUGGAAAUCUATT-3′; CEP63 Oligo, 5′-GGCUCUGGCUGAACAAUCATT-3′. Target sequence of specific gRNA used against CEP152 in stable cell line generation was 5’-CTGAGCAATTGGAGATGAGC-3’. All siRNAs and gRNAs were purchased from Eurofins Genomics, Ebersberg, Germany, at high-performance liquid chromatography purity.

### Plasmid constructions and molecular cloning

Full-length human CEP152 with gRNA resistance (5’-CAGAACAGTTAGAAATGAGT-3’) in pAID 1.1C-T2A-Bsr (Addgene) and CEP152^D43/44A^ in pEGFP-C1 were synthesized by GenScript. pEGFP-C1 (Clonetech), pEGFP-C1-CEP152 WT, pEGFP-C1-CEP152 (1-512aa), pEGFP-C1-CEP152 (508-1710aa), pCMV-3Tag1A-Flag (Agilent Technologies), pCMV-3Tag1A-Flag-PLK4, pCMV-3Tag1A-Flag-PLK4^K41R^, and pCMV-Myc-PLK4 were provided by the lab (Cizmecioglu et al., 2010). pEGFP-C1-CEP192 was provided by Kyung Lee (Kim et al., 2013). CEP152 gRNA sense and antisense were dimerized and cloned into the pSpCas9(BB)-2A-GFP (pX458, Addgene) plasmid by using the provided protocol established by (Ran et al., 2013) available on the Addgene webpage. pGEX-4T1-CEP152 (1-152) was also provided by the lab for recombinant protein expression. Clones were analyzed by sequencing.

## Recombinant protein expression

Human GST-CEP152(1-512aa) was expressed in *Escherichia coli* BL21-Rosetta and purified under denaturing conditions by single-step affinity chromatography using glutathione-agarose according to the instructions of the manufacturer. CEP152 (1-512aa) was dialyzed against PBS, pH 7.0, and concentrated with Vivaspin MWCO 12-15 kDa.

### *In vitro* kinase assay

For *in vitro* kinase assay with recombinant Zz-PLK4-His, GST-CEP152 was purified as described above. 1 μg of Zz-PLK4-His protein was incubated with 2 μg of GST-CEP152 or an equivalent molar amount of GST in kinase buffer (50 mM Tris pH 7.5, 10 μM MgCl_2_, 10 μM MnCl_2_, 1 mM DTT) with or without ATP (0.5mM) for 20 min at 30°C. Reactions were stopped by adding sample buffer and heating at 95°C for 5 min. Samples were analyzed by Western blotting. For the analysis of PLK4 phosphorylation, phospho-specific p-S305 PLK4 antibody was used (Park et al., 2014).

### Cell lysis, co-immunoprecipitation and Western blot analysis

Cell lysates were prepared as described previously (Hanle-Kreidler et al., 2022). Cells were harvested and washed with PBS. For Western blot analysis cells were lysed in RIPA buffer (50 mM Tris–HCl, pH 7.4, 1% NP-40, 0.5% sodium deoxycholate, 0.1% SDS, 150 mM NaCl, 2 mM EDTA, and 50 mM NaF) and for co-immunoprecipitation experiments transfected HEK293T cells were lysed in NP40 lysis buffer (40 mM Tris, pH 7.5, 150 mM NaCl, 0.5% NP40, 5 mM EDTA, 10 mM β-glycerophosphate, 5 mM NaF, 1 mM dithiothreitol, 0.1 mM Na_3_VO_4_, and protease inhibitors). Anti-Flag M2 Affinity Gel was used to pull-down Flag-tagged proteins. Beads were washed 3 times with TBS (50 mM Trizma-HCl, with 150 mM NaCl, pH 7.4), once with 0.1 M glycine HCl, pH 3.5 and again 3 times with TBS (50 mM Trizma-HCl, with 150 mM NaCl, pH 7.4). Cell lysates and beads were incubated on a rotating wheel overnight at 4°C. Beads were washed three times with NP40 buffer and proteins were eluted with 3X flag peptide with 250 ng/μl concentration in NP40 buffer for 30 min.

Samples were run on SDS-PAGE, and proteins were transferred onto a nitrocellulose membrane by using Bio-Rad Trans-Blot turbo transfer system. All the membranes were blocked with 5% Skim milk in PBS-T. Blocked membranes were incubated with determined primary antibody solutions which were prepared in 5% Skim milk in PBS-T overnight. After primary antibodies were collected, membranes were washed three times with PBS-T and incubated with HRP-conjugated mouse or rabbit secondary antibodies for 1 h at room temperature.

### Expansion microscopy

The ultrastructure expansion microscopy (U-ExM) protocol was adapted from previously described protocol by (Gambarotto et al., 2021). U2OS CEP152-AID cells were seeded on 12mm coverslips. After the treatments were completed, pre-extraction was performed prior to MeOH fixation. Fixed samples were incubated with formaldehyde/acrylamide mixture for 4 h at 37°C. Gel polymerization, protein denaturation and first round of gel expansion were performed as described by (Gambarotto et al., 2021). To label the proteins with primary antibody, gels were incubated with primary antibody solution in 2% BSA in PBS for 2 days at 4°C. The labelled gels were stained with the secondary antibodies for 2-3 h at 37°C and the second round of expansion was performed. Expanded gels were fixed in a 35 mm glass bottom imaging chamber coated with Poly-D-Lysine. Images were acquired as Z-stacks and were deconvolved by using either Zeiss or Huygens deconvolution software.

### Immunofluorescence of fixed specimens

For indirect IF, HeLa and U2OS cells were treated as described previously in this paper. Coverslips were incubated in BRB80 pre-extraction buffer (80mM PIPES pH 6.8, 1mM MgCl_2_, 1 mM EGTA and 0.1% TritonX-100) for 30 s and then fixed with ice-cold MeOH for 10 min at -20°C. Cells were blocked in blocking buffer (3% BSA, 0.1% Triton X-100 and 0.2% NaN_3_ in PBS) for 20 min. Both primary and secondary antibodies were prepared in blocking solution. Coverslips were incubated with primary antibodies for 1 h at room temperature and washed with PBS for 3 times. After 1 h of secondary antibody incubation, the coverslips were washed again with PBS for 3 times and were mounted onto glass slides with ProLong Gold with DAPI (Molecular Probes). For cell imaging in both IF and U-ExM, the Zeiss motorized inverted Observer.Z1 microscope was used, containing mercury arc burner HXP 120 C and LED module Colibri. Filter combinations: DAPI, 4′,6-diamidino-2-phenylindole (49), GFP (38 HE) DsRed (43 HE), and HXP120 for Far-Red, and with the detector gray-scale charge-coupled device (CCD) camera AxioCam MRm system and 40x or 63 ×/1.4 Oil Pln Apo DICII objective. Zeiss Apotome confocal imaging device was used during z-stack imaging. For the signal intensity analysis, all images in the same set of experiments were acquired with the same settings (exposure time, Z-stack size and Z-stack number) and maximum projection images were used during the analysis. Centrosomal signal intensity of PLK4, p-S305 PLK4 or CEP152 was quantified by randomly choosing cells by drawing a 3.5 μm^2^ circle around the centrosome. The same circle was used to measure the background signal within the cells and this signal was subtracted from the measured signal from the centrosome.

### Statistical analysis

GraphPad Prism, version 10 (GraphPad Software, Inc.) was used to perform a statistical analysis of the data collected from at least three independent experiments. The data was represented as either individual values or the mean ± SD. To perform a statistical analysis of fold change data, all values were normalized to a control group and a logarithmic transformation was applied to obtain normally distributed data. Logarithmically transformed values were further analyzed by a one-sample, two-tailed t test against the mean of the control group, which was set to 1.0, or by a paired, two tailed t test for comparisons among the test groups to determine the significance. If data was normally distributed and not normalized to a control group, an unpaired, two-tailed t test with Welch’s correction was used for the statistical analysis. P-values below 0.05 were considered statistically significant (ns P > 0.05, *P < 0.05, **P < 0.01, ***P < 0.001, and ****P < 0.0001).

## ACKNOWLEDEGMENTS

We thank Kyung Lee, NIH, USA for reagents, Jadranka Loncarek, NIH, USA for sharing their detailed protocol to generate AID-cell lines, Holger Lorenz, University of Heidelberg, for help with the deconvolution software, and to Alexandra Turi da Fonte Dias for comments on the manuscript. Sophia Grossi-Mayer is thanked for proofreading.

## SUPPLEMENTAL FIGURE LEGENDS

**Figure S1: Fast degradation of CEP152 fused to an AID in U2OS cells. (A)** Sequencing result confirming the inactivation of CEP152 alleles in U2OS CEP152-AID cells by a deletion mutation. **(B)** U2OS CEP152-AID or U2OS WT cells were incubated or not with IAA for 2 h. Cells were then harvested for Western blot analysis. **(C)** U2OS CEP152-AID cells treated with or without IAA for indicated time points. Cells were lysed and analyzed in Western blot analysis. **(D)** U2OS CEP152-AID cells were treated with or without IAA for 2 h, and IAA washed-out for indicated time points and lysed for immunostaining. **(E)** Immunofluorescence staining of U2OS CEP152-AID cells shows fast degradation of CEP152 upon 2 h IAA treatment. Scale bar: 1 μm. CEP152 signal around centrosome upon 2 h IAA treatment is quantified and the background signal was subtracted. Values were normalized to the non-treated control and individual values were presented in the graph with ± SD. n = 2, * < 0.01. **(F)** U2OS CEP152-AID cells treated with IAA for 2 h prior to analysis by U-ExM. Gels were stained with acetylated tubulin and the length and the diameter of centrioles were measured. The numbers obtained were divided by expansion factor of the gels to obtain biological values. Scale bar: 2 μm, biological scale: 0.6 μm. Graphs represent the quantification of centriole length and diameter, n = 2, ns > 0.05.

**Figure S2: CEP152 collaborates with CEP63 for PLK4 recruitment to the centrosome. (A)** HeLa CEP152-AID cells transfected with control and CEP63 siRNAs and treated with or without IAA for 2 h. Cells were fixed for U-ExM analysis and then stained with acetylated tubulin to mark centrioles. Scale bar: 2 μm, biological scale: 0.5 μm. Graphs represent the quantification of centriole length and diameter after the values obtained from measurement were divided by expansion factor of the gels to determine biological values n = 2, ns > 0.05. **(B)** U2OS CEP152-AID cells transfected with control (siGL2) and CEP63 siRNAs and treated with or without IAA for 2 h. Fixed samples were stained with antibodies against PLK4 and y-tubulin. Images show the change in PLK4 levels at the centrosome. Scale bar: 1 μm. PLK4 signal was normalized to the non-treated control and values were shown in the graph with ± SD n = 2. * = 0.05, scale bar: 1 μm.

**Figure S3: Complete removal of CEP152 does not change the level of PLK4 at the centrosome but decreases PLK4 activation. (A)** U-ExM images of HeLa CEP152-AID cells transfected with siGL2 and siCEP152 for 48 h and treated with - / + IAA for 2 h. Graphs represent the quantification of centriole length and diameter n = 1, ns > 0.05. **(B)** U2OS CEP152-AID cells were transfected with the indicated siRNAs for 48 h and treated with - / + IAA for 2 h prior to fixation. The samples were immunostained. Scale bar: 1 μm. p-S305 PLK4 and PLK4 signals were quantified in cells with no detectable CEP152 signal at the centrosome and normalized to the control after background signal subtraction. Graphs show individual values of control normalized p-S305 PLK4 signal. n = 2, ns > 0.05, * = 0.05, ** < 0.01. **(C)** HeLa CEP152-AID cells were transfected with control and CEP152 siRNAs and treated with - / + IAA for 2 h. Fixed coverslips were stained with either Ninein (mother centriole marker) on the left panel or Centrobin (daughter centriole marker) on the right together with p-S305 PLK4 and centrin. Scale bar: 1 μm. n = 3, * = 0.05, ** < 0.01.

