## Supplementary figures and images for "Binding of CEP152 to PLK4 stimulates kinase activity to promote centriole assembly"

### Supplemental Figure 1

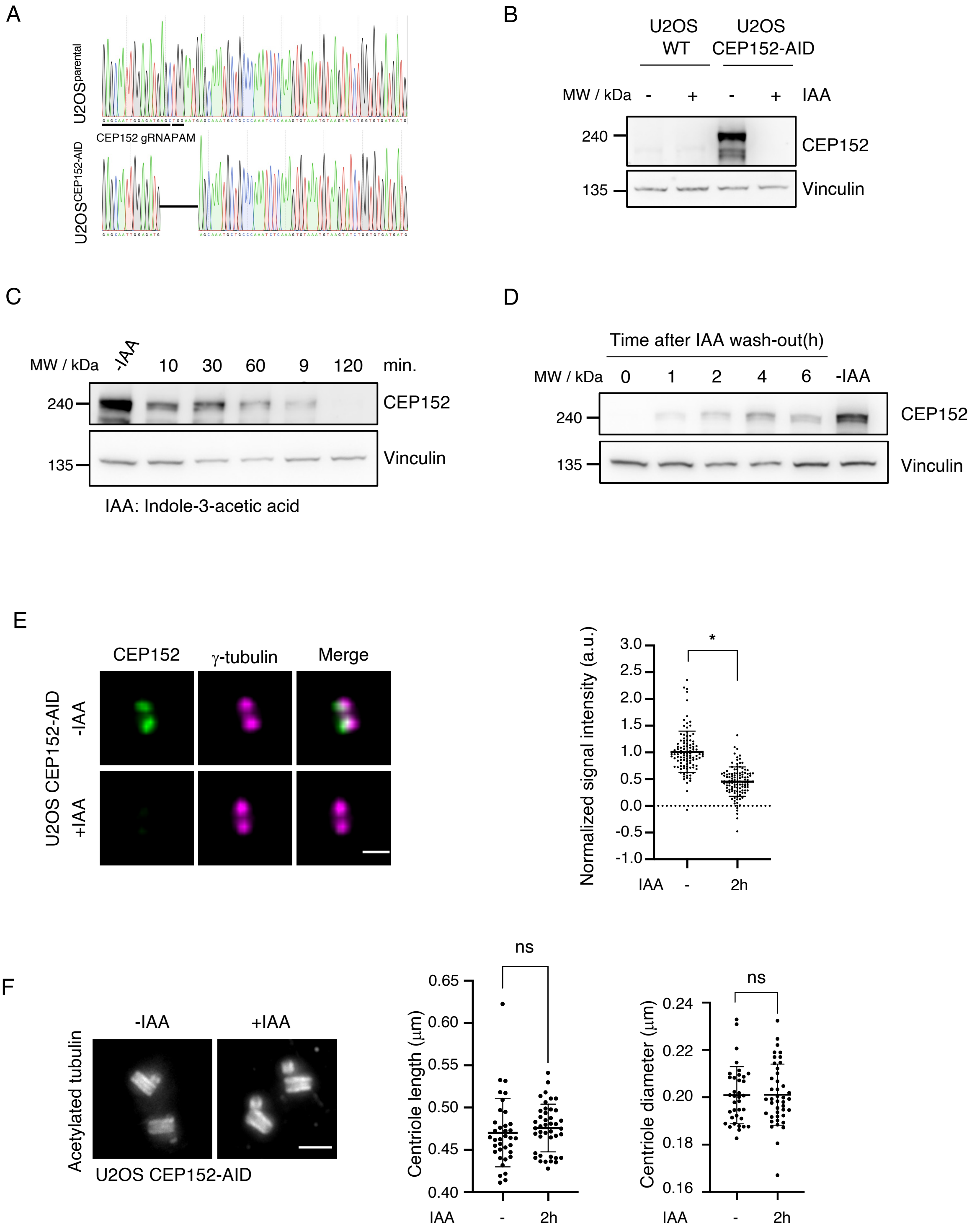

### Supplemental Figure 2

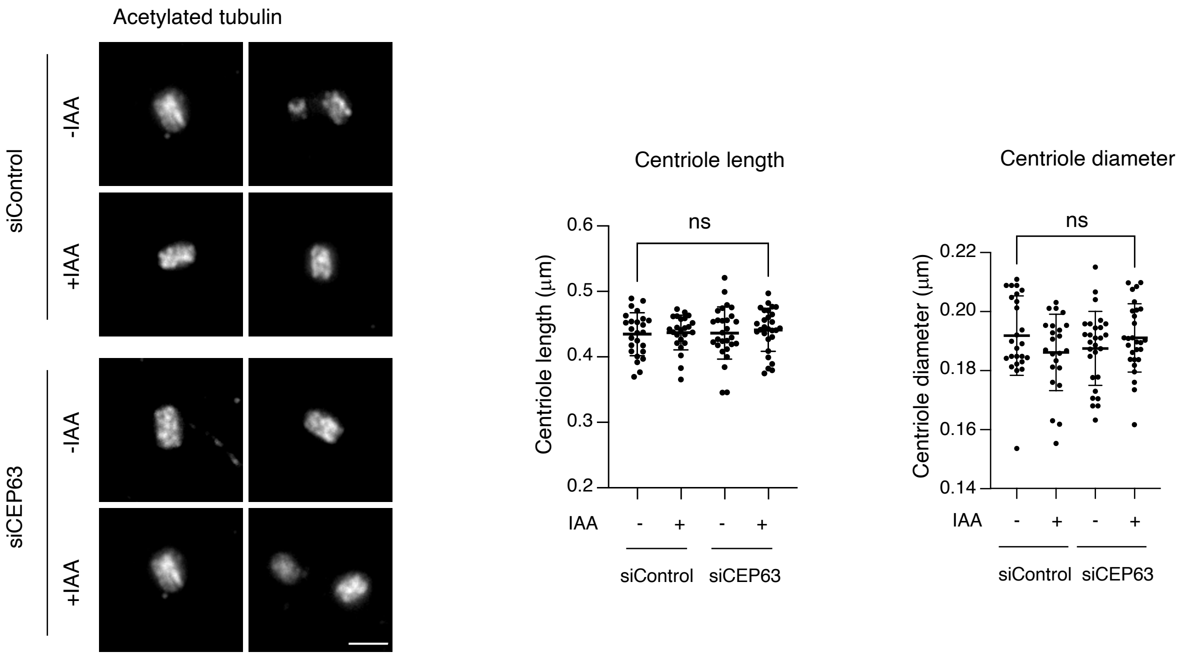

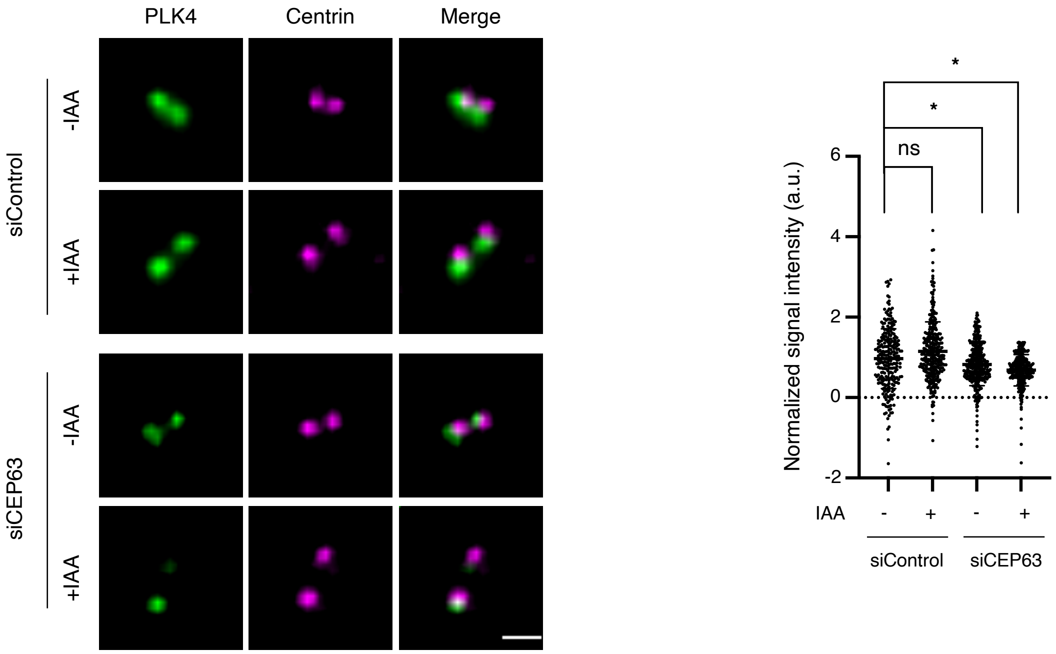

### Supplemental Figure 3

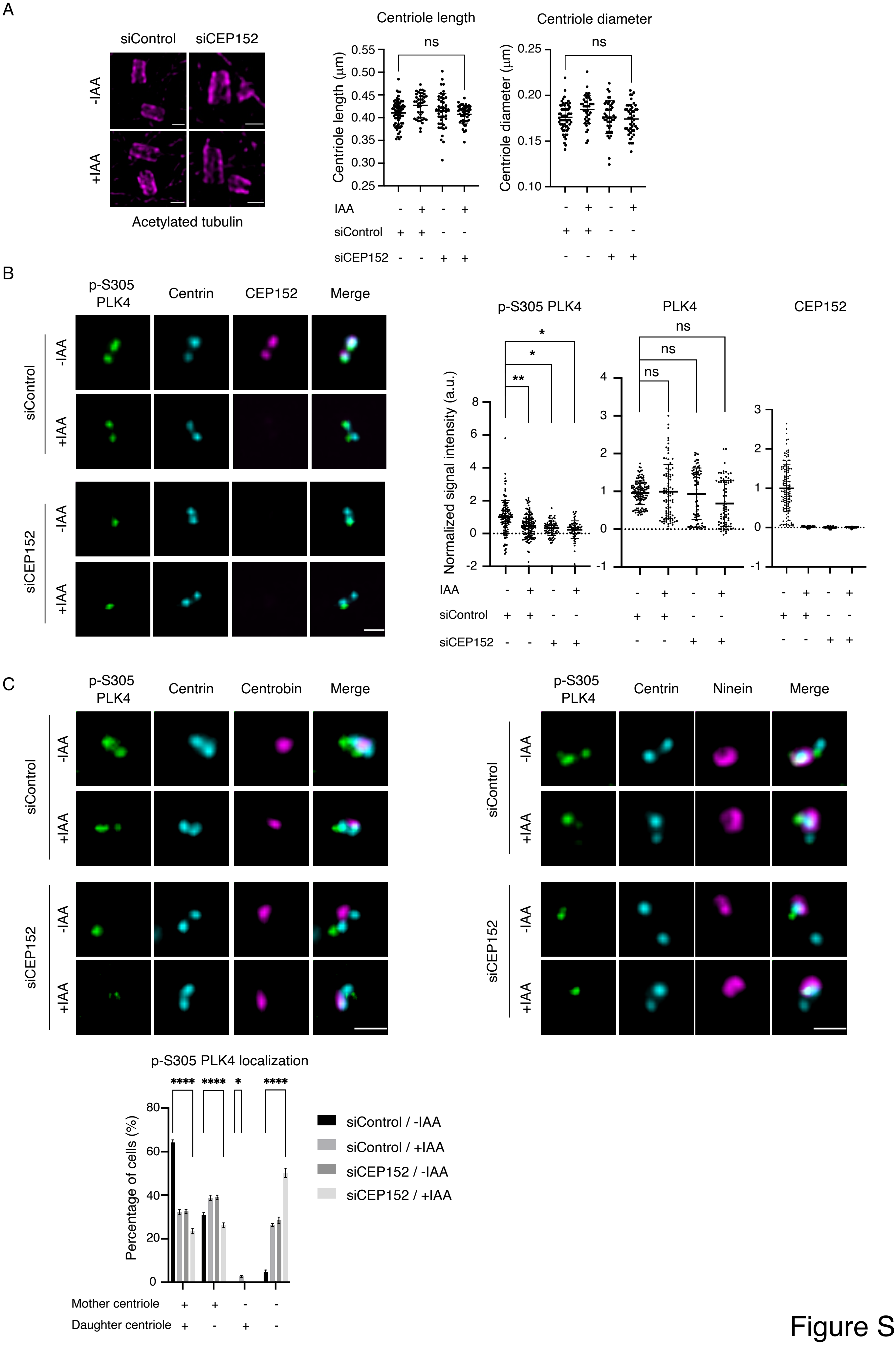
